# Spatial transcriptome sequencing revealed spatial trajectory in the Non-Small Cell Lung Carcinoma

**DOI:** 10.1101/2021.04.26.441394

**Authors:** Li Zhang, Shengqiang Mao, Menglin Yao, Ningning Chao, Ying Yang, Yinyun Ni, Tingting Song, Zhiqiang Liu, Yongfeng Yang, Weimin Li

## Abstract

Deepening understanding in the heterogeneity of tumors is critical for clinical treatment. Here we investigate tissue-wide gene expression heterogeneity throughout a multifocal lung cancer using the spatial transcriptomics (ST) technology. We identified gene expression gradients in stroma adjacent to tumor regions that allow for re-understanding of the tumor micro-environment. The establishment of these profiles was the first step towards an unbiased view of lung cancer and can serve as a dictionary for future studies. Tumor subclones were detected by ST technology in our research, while we contrast the EMT ability among in subclones which inferred the possible trajectory of tumor metastasis and invasion, which was confirmed by constructing the pseudo-time model of spatial transition within subclones. Together, these results uncovered lung cancer spatial heterogeneity, highlight potential tumor micro-environment differences and spatial evolution trajectory, and served as a resource for further investigation of tumor microenvironment.

## Introduction

Lung cancer has been the leading cause of cancer death allover the world, accounting for 22.7% of malignant tumor classification. Non-small cell lung cancer (NSCLC) accounts for more than 80% of all lung cancers, with two major histological subtypes are lung adenocarcinoma (40%~50%, LUAD) and lung squamous cell carcinoma (30%, LUSC) (Sakashita S et al., 2014). As we know gene expression profiles, epigenetic and tumor origin are important in influencing cellular programs and lead to disparate disease pathogenesis of NSCLC (Herbstet al., 2018; Shiet al., 2019). In addition, with LUSC exhibits faster progression rate than LUAD (Chen et al.,2017, Xiang et al., 2020). To understanding of the tumor micro-environment and mechanism of cancer metastasis is important for future treatment of NSCLC.

In the past decades, transcriptomic studies have revolutionized cancer treatment and diagnosis. In recent years, by using single-cell RNA-Seq (scRNA-Seq), the gene expression heterogeneity of NSCLC has been documented at the level of individual cell level. Previous studies investigating lung cancer have primarily focused on the landscape of immune and infiltrating cell populations (Guo et al., 2018) or have specifically identified novel cell subtypes and altered pathways (Lambrechts *et al*., 2018). However, scRNA-seq studies often suffer from lack of tissue morphology transcriptome expression and loss of spatial information during tissue dissociation. In recent years, the technology could simultaneous determine of the locations of dozens of cell types within the tumor, which is important for understanding the components of the tumor micro-environment and tumor-stroma crosstalk (Angelo et al., 2014; Keren et al., 2018; Salmén et al., 2018; Thrane et al., 2018). Spatial transcriptomics (ST) presents a solution to this dilemma, by providing spatially resolved and transcriptome-wide expression information. For example, the success of immunotherapy in tumor treatment stems from its ability to interfere with interactions between cancer and certain immune cells (Farkona, S. et al.,2016 and Ståhl, et al.,2016).

Tumor have genetic instability and highly variable that leading to the different tumor cells, that is tumor subclones with different mutation and manifestation, tumor subclones can undergo “linear” or “branch” evolution under different selective pressures. Subclones in the original metastasis play an important role in tumor development. Therefore, the evolution trajectory of subclones is important for tumor evaluation and clinical guidance. scRNA-seq has been used to inference in vitro cell lineage trajectories widely, by identifying trajectories that connect cells based on similarity in gene expression, one can gain insights into lineage relationships and developmental trajectories (Saelens, et al., 2019). Traditionally, most statistical models to study cell progressions for instance, across cell differentiation or cancer evolution, have not considered spatial structure. There is not yet a survey that reconstructed biological trajectories within a tissue. In this work, we adapt spatial pseudo-time method to understand how the tumor invasion (infiltration) and metastasis on the intra-tumoral (Pham, et al., 2020).

Here, in this study, we used ST technology to survey the spatial patterns of gene expression and cell types in 4 LUAD samples and 8 LUSC samples collected from twelve individuals. Intra-and inter-patient heterogeneity was examined using different methods, including expression-based clustering and morphological analysis. We found that the LUSC more invasive the surrounding and stromal and higher intratumoral heterogeneity in the spatial microenvironment, more suitable as a model of microenvironment and tumor subclones evolution trajectory. We identified the subclones within the tumor and found that EMT-related pathways have significant differences in tumor metastasis, implying that they may have undergone different differentiation processes. Therefore, we used the spatial-pseudo-time method to construct the possible spatial evolution Trajectory between the subclones. Our finding firstly try to model the tumor metastasis trajectory on the slice by ST data, which will help us to more clearly observe the tumor spread and metastasis, and propose better diagnosis and treatment in clinical.

## Results

### Overview of study the lung cancer spatial data

We performed Spatial Transcriptomics on lung tumor tissues collected from 12 individuals presenting for surgical resection using the 10x Genomics Visium platform (Figures S1, Methods).The collected sections of twelve patients (LUAD 4 samples & LUSC 8samples) were then subjected to spatial transcriptomics using circular spatial spots with a diameter of 100μm, each section was carefully examined by an experienced pathologist, who annotated the different morphological regions in the section, including tumor, stromal, immune cell et al. These annotations either served as a reference for subsequent unsupervised analysis or were used directly to pick spatial spots from regions of interest. We conducted ST library for each sample, mainly focusing on tumor region and the important tissue regions that we interested in for the analysis of spatial micro-environmental gene expression, identification cells type, the reconstruction of cell trajectories, cell-cell interactions within a morphologically-intact tissue section (Figure.1a).

**Figure 1.**
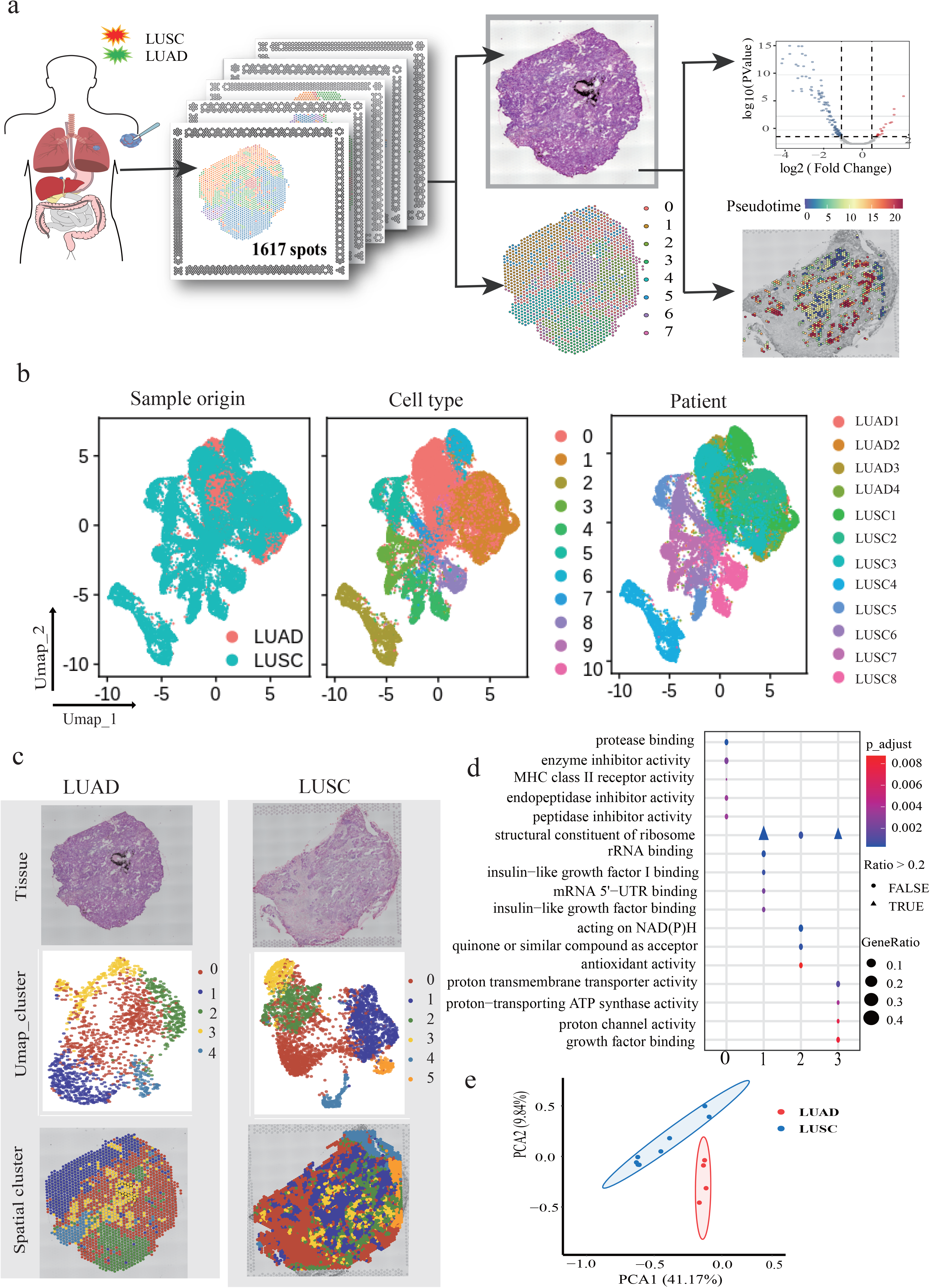
Overview of the whole study. a: Workflow of Spatial Transcriptomic lung cancer, The LUAD) and LUSC of non-small lung cancer as experimental materials. 1: Location of sections used in this study and annotations made by the experienced pathologist. 2 Spatial microarrays have 1617 spatially barcoded spots. The ST procedure yields matrices with read counts for every gene in every spot. 3: Each tumor tissue section can accurately reflect the spatial specific expression characteristics of genes through spatial transcriptome data.4: Finally, select the tissue regions we are interested in for the analysis of spatial micro-environmental gene expression and the identification cell type, the reconstruction of cell trajectories, the study of cell-cell interactions within a morphologically-intact tissue section. b: Summary of the sample origins. (b) UMAP visualization of combined pixels from LUAD samples and LUSC samples using Seurat package. Left: UMAP labelled by sample names; In the middle UMAP of the 11 samples profiled here, with each cell color-coded for (left to right), the fraction of cells originating from the 4 LUAD patients and 8 LUSC patients; right: UMAP labelled by cluster numbers. Totally 10 clusters were identified. c: The two samples contain LUAD1 and LUSC1 was selected to show: Tissue morphology, anatomical annotation, and spatial mapping of the Seurat clusters in middle, the pathologist’s annotations are well agreement with the results of cell clustering. d: GO enrichment analysis of all clusters (0-5). e: Comparison of “pseudo bulk” data between LUAD and LUSC reference data.

Integrated clustering analysis of ST from all samples using Seurat (*v4.0*) revealed 10 distinct clusters, the spot type between LUAD and LUSC had a great difference (Figure.1b). Considered of tumor heterogeneity and individual differences of different patients, we next analyzed each section individually to ensure the accuracy of result. To better understand the distribution of these cell types that have been divided spatial distribution, we mapped the clusters back to their spatial location identified spatially distinct patterns that could match the anatomical annotation (Figure.1c & Figures S1). The example of LUAD_1, respectively cluster 0 high expression of *SFTPC, STTPD* which mainly represents the AT2 cells. Cluster 1 major covers the secretory club cells in the tissues which specific expressed *SLPL, SCGB3A1*. Cluster 2 is the tumor tissues, cancer related cells is largely shown as cluster 2, *SCD, SFTA2, DMBT1* which are tumor related marker higher expression than other clusters. Cluster 3 identified specific expression genes of *AOPC1, CHIT1*, Cluster 4 was considered tumor stroma tissues mainly represented by fibroblasts. We further conducted GO analysis (Figure.1d) for each cluster, and the GO pathways matched well with the anatomical annotation and further explain the biological process of the subgroup. We further compare of “pseudo bulk” data between LUAD and LUSC reference data, within the same cancer type have the well compatibility (Figure.1e), the significant difference between the LAUD and LUSC indicated that they maybe have different spatial transcription expression patterns.

### Transcriptome heterogeneity in the spatial TME between LUAD and LUSC

Bulk RNA-seq has proved that there are great distinction in gene expression profiles between LUAD and LUSC homogeneous expression level in the whole tissue (Michael A. et al., 2019). We sought to investigate tumor spatial microenvironment differences in gene expression between the tumor and the adjacent stromal tissues, in order to understand the difference more accurately between the expression profiles of tumor cells and the surrounding stromal cells. We distinguished tumor and adjacent tissue and thereby provided insight into gene expression variations during the progression of lung cancer. The pathologist annotated the region by HE stained sections (Figure 2a). Then, extract the barcodes cells in the corresponding area, and then analyzed the different gene expression between these areas. We found that genes function related to cancer malignant and are highly expressed in LUAD_1 (*SLPI, SCGB3A1, SCGB3A2, MS4A15, NR4A1*) (Toshihiro et al.,2008; Susan D. et al.,2002; Sun et al.,2019), related to tumor metastasis and tumor cell proliferation such as *SPP1*, *GSTA1, MAL2, MGST1* are highly expressed in LUSC_1, LUSC_2, LUSC_4 samples (Figure 2a &2d) (Wei et al.,2020; Pan et al., 2014; Eguchi, et al., 2013). The function of DEGs revealed the high activity of peptidase in LUSC in metabolism pathways (Figure 2b&2e), cell-substrate adhesion is different in the tumor microenvironment of the biological process. However, no evaluation with spatial resolution of lung cancer and cancer stroma tissue has been performed to date. We therefore aimed for discover differences between the tumor micro-environment on same section.

**Figure 2.**
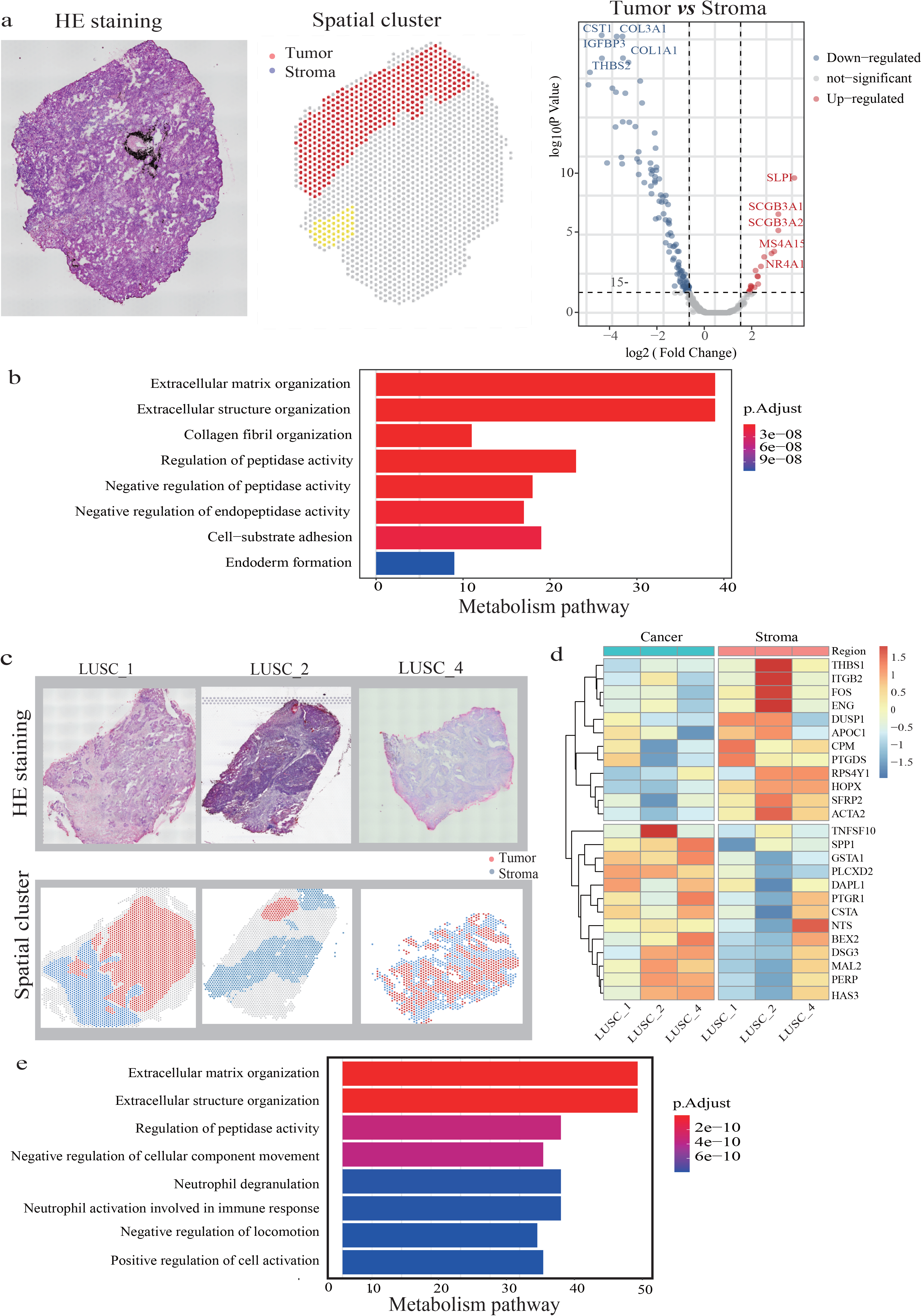
Spatial comparison of tumor stroma and tumor transcriptomes between the lung cancer. Tissue sample: Tissue sample area comprising spots taken for normalization of ST counts, within this area spots are chosen as tumor stroma and cancer. Choice of spots is based on the pathologist’s annotation and the activity of the factor cancer and reactive stroma, Volcano plot of significantly differentially expressed genes between periphery and center. b, Box plots showing expression levels of noteworthy genes significantly upregulated in either tumor stroma or the cancer enriched pathways for significantly (p < 0.05) differentially expressed genes in cancer and stroma. P-values per gene were calculated with two sample t-test. **a & b**: The different tumor stroma and tumor transcriptomes expression in the LUAD sample (LUAD1) and the different expression pathways are shown in the picture. **c& d**: Different spatial transcriptomes expression in the tumor micro-environment in the LUSC sample. (LUSC1) and the different expression pathways are shown in the picture.

To compare the tumor microenvironment of two lung cancer subtypes of NSCLC, we significantly enriched more pathways related to epithelial-to-mesenchymal transition in LUSC in Spatial microenvironment (Figure 2e). The DEGs also showed results of experiments have been proven that up-regulation of *SPP1* enhanced cell viability and promoted tumor cell proliferation and *MAL2, MGST1* promotes proliferation, migration, and invasion through regulating epithelial-mesenchymal transition in breast cancer cell lines, we speculate them may perform equivalent function in LUSC.

Taken together, the LUSC exhibited more invasive ability which can be metastasized direct surrounding and lymph node metastasis, less distant metastasis, rich immune microenvironment and cell communication, which suggested that LUSC was more suitable as a model of tumor spatial microenvironment and tumor subclones evolution trajectory than LUAD in the NSCLC.

### Pattern of transient gene expression along spatial trajectories

The phenomenon that a malignant tumor become more likely to aggressive as they grow is called tumor evolutional metastasizes. The main surface is the tumor spreading, metastasis and invasion of tumor stromal tissue. To further explore the changes in gene expression along the path of tumor invasion, we adapted the spatial trajectory analysis with SPATA (Jan Kueckelhaus, et al.,2020), which introduced a new approach to find gene expression pattern, we analyze and visualize differently expressed genes in a spatial lung cancer section. While the classic differential gene expression analyzes the differences between different groups, as the whole it neglects changes of expression levels that can only be seen while maintaining the spatial dimensions.

Pathology selected section that was in the stage of tumor spreading (Figure 3a), which considered to be the model for tumor metastasis (LUSC_3). Firstly, we signed the invasion direction of the tumor and chose the trajectories, Visualization of continuous features along a trajectory that found the proportion of cancer spot gradually decrease and stroma spot rise, immune-related cells have no significant changes (Figure 3b) Visualization of discrete gene features along a trajectory, we found associated with promotes tumor proliferation like *DSG2* and *SPRR3* gene modulates cell proliferation, invasion and cell apoptosis in NSCLC were decreased, tumor progression (*BGN*) and metastatic growth (*POSTN*) function gradually enhanced. To find variables that follow a certain trend along the trajectory, we created a set of mathematical models to model the gene expression of single genes which represent defined biological behaviors, including linear, logarithmic or gradient ascending/descending expression pattern, one-, or multiple peak expression. We focused on genes whose expression increase or decrease in expression along the trajectory. The eight patterns models were observed (Figure 3b) and we found that related to the oxidative phosphorylation and ATP synthesis coupled, electron transport ATP metabolic process pathways decreased along the trajectory of the tumor to stroma, while extracellular matrix organization, Immune response, antigen processing and presentation were enhanced in tumor stroma. The increased energy requirement of tumors mainly came from glycolysis and aerobic oxidation of mitochondria for tumor proliferation, which tumor stroma contained macrophages, T cells, B cells or other immune cells, which participate in immune response, affect tumor microenvironment, and regulate tumor development and metastasis. (Figure 3b & Figure 3e)

**Figure 3.**
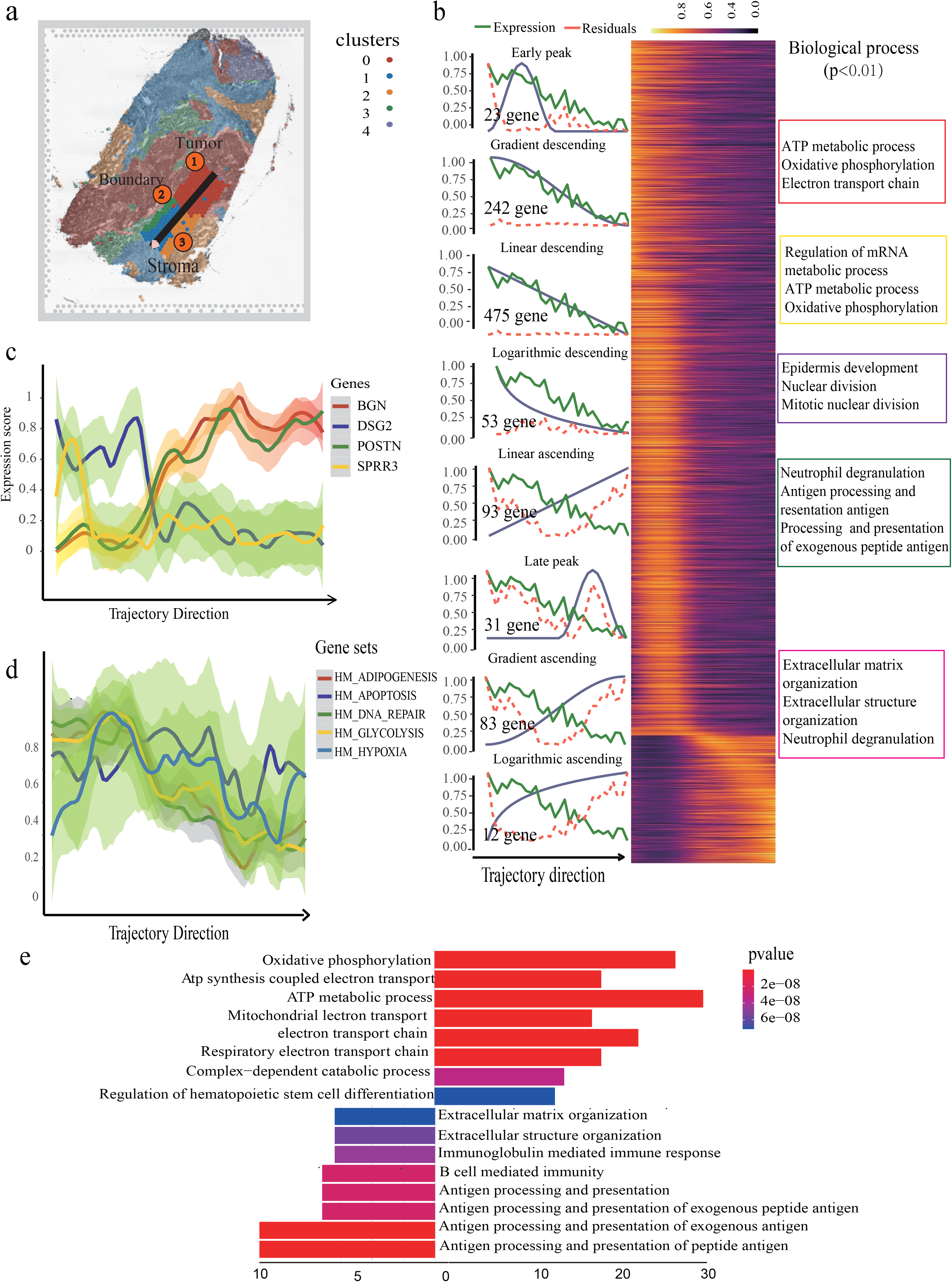
Pattern of transient gene expression along spatial trajectories. (a) Draw the trajectory interactively by clicking on the locations where you want the start and endpoints to be. A: For LUSC_3 we specify a trajectory on a section, B Visualization of discrete features along a trajectory. (b) Changes in gene expression along a spatial trajectory (BGN, DSG2, POSTN, SPRR3). (3) Visualization of continuous features along a trajectory, gene-set of biological pathways, Such as (HALLMARK _ADIPOGENESIS, HALLMARK _APOPTOSIS, HALLMARK _DNA_REPAIR, HALLMARK _GLYCOLYSIS, HALLMARK _HYPOXIA). (4) Visualization of the fitting results. (5) To get a summary of your trajectory assessment use examine TrajectoryAssessment() which displays the distribution of all areas under the curve measured(left). Heatmaps to display dynamic expression changes along trajectories (right).

### The heterogeneity of tumor subclones spatial expression

Intra-tumoral is highly heterogeneous, even the same malignant tumor is composed of different subclones, which can lead to the failure of targeted therapies and drug resistance. We attempted to investigate the heterogeneity of subclones by using ST data. Tumor tissues harbored different cell and tissue morphology from normal tissues in the LUSC, ST technology adapt H&E Staining can distinguish the degree of malignancy of cancer cell. The pathologists help us to detected tumor region spots of each section. Thus, copy number variation was a major feature of tumor cells. We used inferCNV software to calculate copy number analysis of spatial transcriptome (Patel et al., 2014). Stroma cells were selected as the reference, we determined that all the subcluster are malignant cells and subcluster all contain different degrees of gain and loss of copy number, which confirmed the result of histological identification of cancer region by the pathologist (Figures S2). Re-clustering these cancer cells revealed 4 clusters. The genes *PI3*+, *KRT17*+ highly expressed in Cluster 0corresponds to basal cell and three clusters to unknown cells (cluster 1; *SPP1*+, *SPARCL1*+), cluster 2; *ACTL6A*+, *NDUFB5*+) and cluster 3; (*KRT5*+ and *KRT15*+). Surprisingly, they occupied a unique region in the section, implying that they might have the functional connection in region. Pathway analyses successfully identified differences between these subgroups of tumor cells; we identify differences in cancer subgroup cells, revealing tumor-associated invasion and metastasis, oxidative phosphorylation, cell proliferation and adaptive inflammatory host defenses, pathways associated with ECM receptor interaction, focal adhesion, and cell adhesion molecules cams.

To further assess the spatial organization of tumor cell populations, we scored spots of the subcluster in each section with EMT-signature genes from Epithelial-mesenchymal transition gene set, which contained 13 mesenchymal marker genes (*VIM, CDH2, FOXC2, SNAI1, SNAI2, TWIST1, FN1, ITGB6, MMP2, MMP3, MMP9, SOX10, GCS*) and three epithelial marker genes (*CDH1, DSP, OCLN*), pan cancer analysis showed that high EMT value of these genes is significantly related to poor survival (Mark D et al., 2015; Jüri et al., 2013; Dev et al., 2018). These MET score high clusters were similar across subcluster among patients (LUSC_1, LUSC_3), which have the significant statistical differences (p =9.3e−06, p < 2.22e−16.). Collectively, these data identify spatial location-specific subclones, which were different of the epithelial-mesenchymal transition level associated with invasion and metastasis. In addition, we used ESTIMATE tools to infer the cellular composition of tumor subcluster and the infiltration of different normal cells (Figures S3). Mainly focus on specific signatures related to stromal cell and immune cell infiltration and tumor purity, the results also showed that the cellular immune composition and the tumor proportion were significantly different in tumor subclones, furthermore tumor purity have opposite trend tumor EMT ability (Figure 3d).

The spatial transcriptome of LUSC could detect tumor subclones, and we found that ability related to tumor migration and invasion were different, provided researchers with an expecting perspective to better understand the tumor heterogeneity. In addition, intra-tumoral heterogeneity was considered to occur during development and tumor metastasis, our experiment also confirmed that, but how did differentiate and originate between these subclones, it is still a concern of researchers.

### The spatial trajectory inference in Intra-tumoral

To discover in vivo processes within tumor, further to answer which cancer cell or clone appeared first or how a cancer evolved, the pseudo-time theory commonly applied in scRNA-seq data analysis, and was designed to detect biological processes that can be inferred from gradient changes in transcriptional states across a tissue. We adapted the pseudo-time trajectory analysis in space using stlearn, which used PAGA trajectory analysis based on tissue-wide SME normalized gene expression data to find connections within subcluster, was applied to study the development trajectory of carcinoma in situ and metastatic carcinoma in breast cancer Pham et al., 2020). In order to model cancer development between the sections, the pseudo-time spatial trajectory algorithm was applied to find the spatial and transcriptional connections among the sub-clusters. In global and local spatial trajectory inference, we adapted pseudo-time spatial trajectory algorithm was applied to find the spatial evolution trajectory and transcriptional connections among the subclones in the LUSC (Figure 5a). The result indicated spatial trajectory direction of tumor cells always starts from cluster1 and direct to other subgroups, showing a divergent state. As well cluster 1 is considered to have low EMT score, which can be explained as the evolution trajectory between subclones is the beginning of the subclones with low metastasis invasiveness and gradually evolves into subclones with strong invasiveness and metastasis ability. By using spatial-pseudotime analysis to detected the subclones progression trajectories, the method allowed us to detect the whole hierarchical tree plot represent for the global spatial trajectory inference among subclones (LUSC_1 & LUSC_3), LUSC_1 have two clades (2,10), the major clade have much branches(0-31,3-34),which was the main inferences of tumor evolution trajectory, LUSC_3 also have similar results a major clade with many branches (4;3-35), reflecting the branched evolution inherent to subclone progression in the LUSC (Figure 5a).

**Figure 4.**
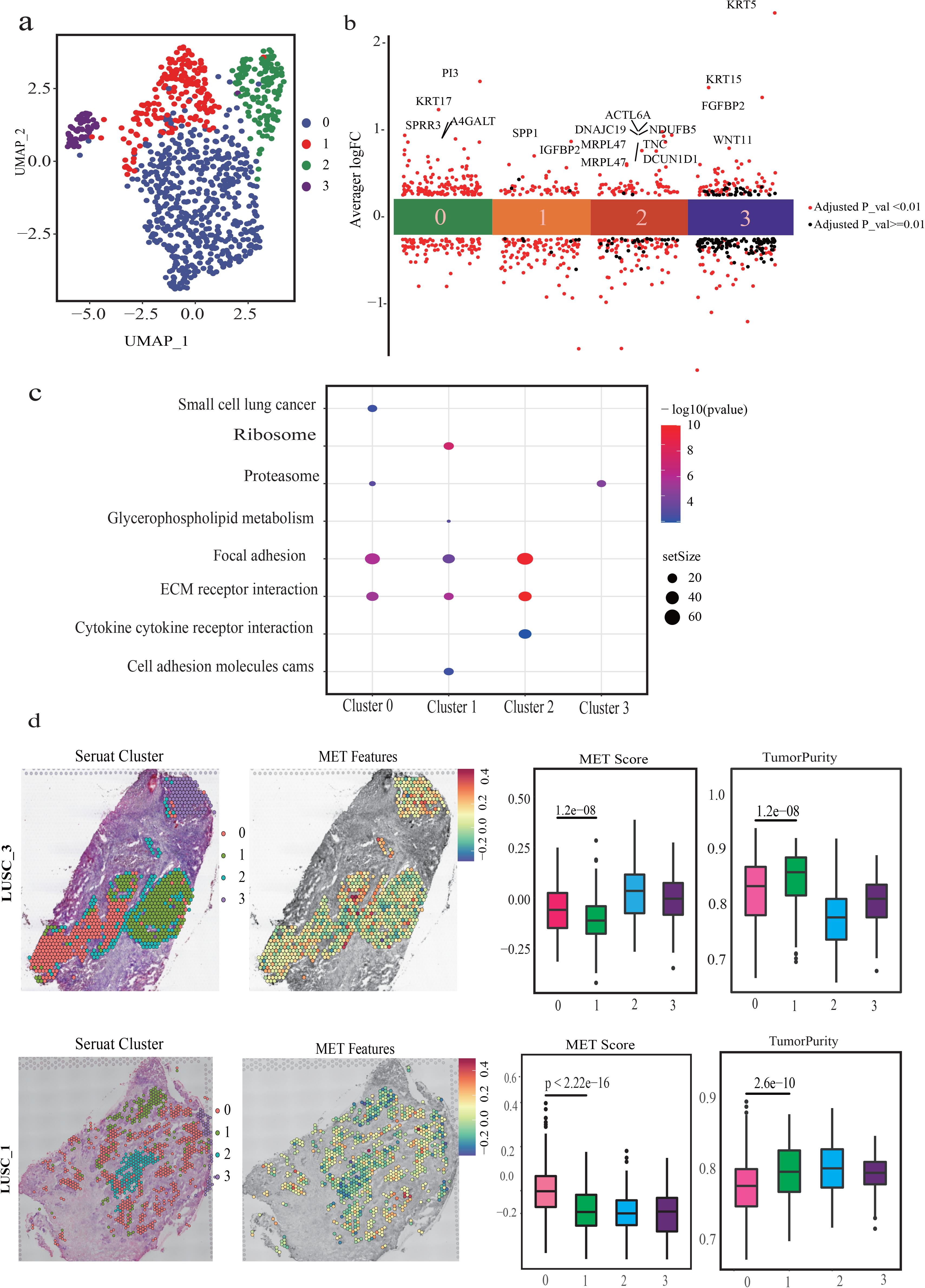
Spots spatial heterogeneity related to tumor formation. a: Re-clustering these cancer cells revealed 4 clusters, cluster 0; *PI3+, KRT17+*) and three clusters to unknown cells (cluster 1; *SPP1*+, *SPARCL1*+), cluster 2; *ACTL6A*+, *NDUFB5*+) and cluster 3; *KRT5*+ *and KRT15*+). c: KEGG pathways of the four subcluster, the results show enriched pathways associated with ECM receptor interaction, Focal adhesion, Cell adhesion molecules cams. d: we adapt the 16 MET gene set proved by the researcher which contain mesenchymal marker genes (*VIM, CDH2, FOXC2, SNAI1, SNAI2, TWIST1, FN1, ITGB6, MMP2, MMP3, MMP9, SOX10, GCS*) and three epithelial marker genes (*CDH1, DSP, OCLN*) and tumor purity which scored each spot of the subclone in each section with EMT-signature genes, T-test was used as a significance test (p<0.001).

**Figure 5.**
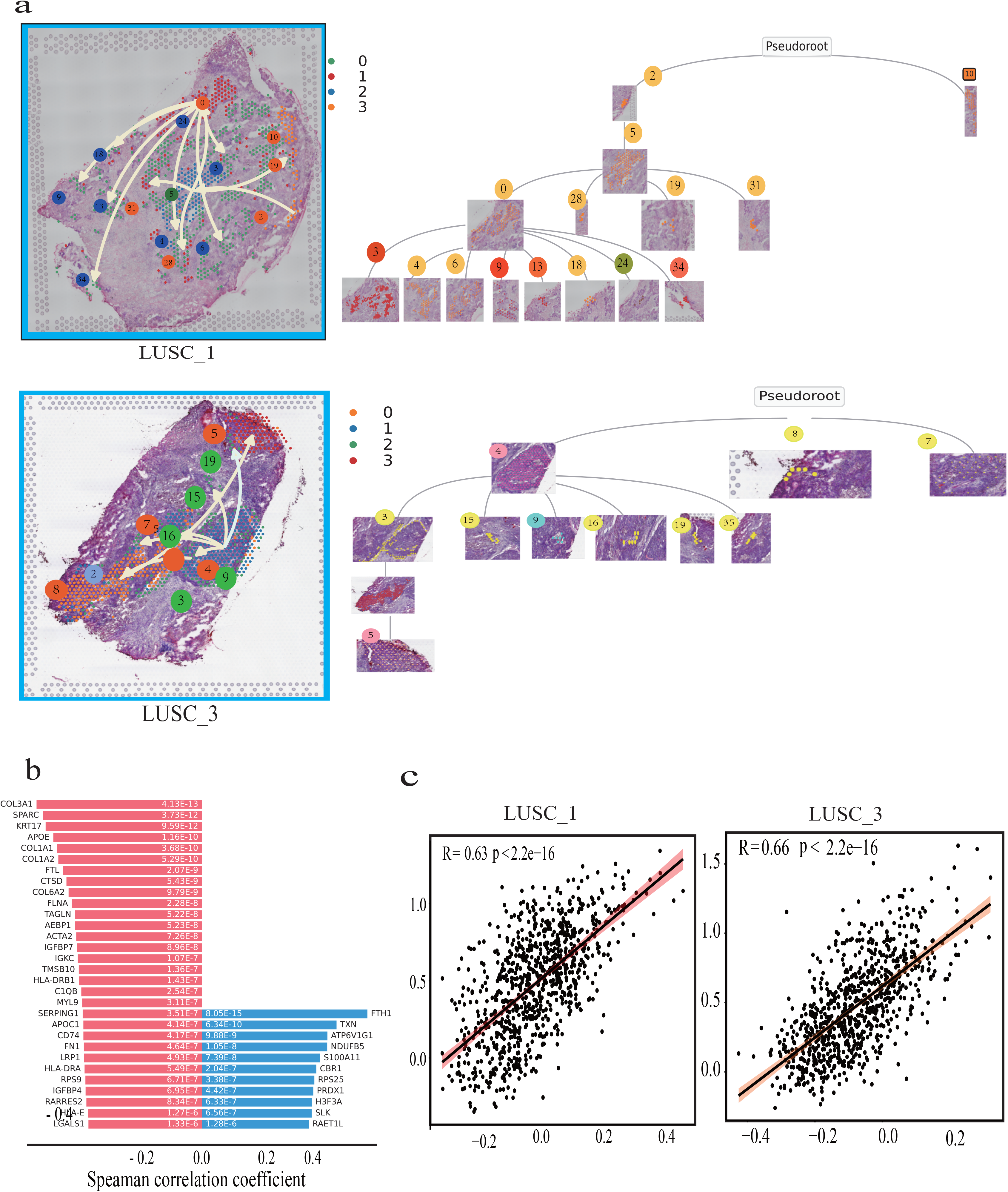
Lung tumor spatial trajectory inference. Spatial pseudo-time-trajectory analysis to discover in vivo processes happening within an intact tissue - for example, to answer which subclones appeared first and cancer evolved. a. The pseudo-time spatial trajectory algorithm was applied to find the spatial evolution trajectory and transcriptional connections among the subclones in the LUSC. b. The trajectory-based differential gene expression which identified transition genes, genes from left side (red) are negatively (down) correlated with the spatial trajectory and genes from right side (blue) are positively correlated with the spatial trajectory c. EMT gene expression and negative spatial trajectory gene expression quantity association analysis in each spot of section.

We calculated the trajectory-based differential gene expression which identified transition genes, genes from left side (red) are negatively (down) correlated with the spatial trajectory and genes from right side (blue) are positively correlated with the spatial trajectory (Figure 5b). In order to further understand the driving factors of subclones spatial trajectory, we combined the gene negatively correlated with the spatial trajectory to a gene set, and scored to tumor related spot, our research found that EMT gene expression and negative spatial trajectory gene have a higher positive related in LUSC_1 R=0.63 and LUSC_2 R=0.66 (Figure 5c).

## Discussion

Here we investigated tissue-wide gene expression heterogeneity in samples from twelve patients diagnosed with lung cancer using spatially resolved transcriptomics. Spatial transcriptome landscape of lung tissue of influenza patients had been recently reported, our study is the first, to implement Visium technology in human lung tumor. Each patient was selected to analyze in a comprehensive method independently that ensure tumor heterogeneity. On average we sequence 20 million unique reads on a tissue slide. Hence, we measure, for the first time, spatial gene expression in lung cancer tissue sections. Further evidence for the accuracy of the spatial gene expression measurements is presented by concordance between gene expression and histological identification within the tissue.

Spatial Transcriptomics technology of lung cancer is the possibility to identify and characterize the tumor stroma or tumor tissue sections in an unbiased way. Unsupervised clustering algorithm based on graph theory, which the spatial transcriptome gene expression profiles may be used to predict further regions of potential cancer, tumor, and angiogenesis region. Pathologists should pay attention to the expression patterns of tumor-related risk genes in specific tumor regions and overall distribution, for clinical research that we suggest spatial gene expression pattern may be useful to gain further understanding of lung tumor in situ or not. Importantly, we report on heterogeneity within one patient as well as heterogeneity between patients. Thus, this study highlights the value of focusing on individual profiling.

Intra-tumor heterogeneity (ITH) contains spatial heterogeneity (different areas of the same tumor) and temporal heterogeneity (different from carcinoma-in-situ and secondary carcinoma), which can lead to low treatment effectiveness and high tumor recurrence. Malignant tumors often proliferate from a single monoclonal malignant cell. During the development, proliferation, clonotypes of tumor differ in growth rate, invasion ability, signal response, and drug sensitivity. This tumor cell population is no longer exactly same as tumor cells, but the “subclone” of heterogeneous tumor cells. EMT refers to epithelial cells that lose cell polarity and junctions, adhesion, increase infiltration and migration and migration capabilities, and become mesenchymal cells. We calculated the EMT score of each tumor subclone to evaluate its ability to metastasize. The result found that different lung cancer subclone cells have independent spatial distribution, and the EMT of each subclone has significant differences, which hints at the trajectory of tumor migration. Besides, we adapt the spatial pseudo time method which integrates spatial location information and gene expression profiles to determine the spatial structure of subclone specific evolution tumor cell states, which modeled cancer evolution trajectory is consistent with the previous result.

Taken together, our findings have revealed that an analysis of lung cancer gene expression in a spatial context dramatically increases the spatial gene expression in the tissue compared to bulk RNA-seq and scRNA-seq. Effective division of tissue sections, and gene expression has remarkable differences on the transcriptome level, which can be used as the clinical evaluation of cancer tissues and microenvironment.

In conclusion, our study established a lung tumor spatial gene expression atlas and extensively investigated the spatial heterogeneity variation between LUAD and LUSC, which could be used as a valuable resource for understanding the unique mechanism of tumor progression in lung cancer subtypes, providing considerable opportunity to advance diagnostic procedures and help design personalized treatments from the time of diagnosis.

## Methods

### Experimental Subject and Method Details

#### Sample Collection

Fresh tumor tissues and adjacent normal tissues were collected from patients with primary lung cancer undergoing surgical resection without neoadjuvant therapy before surgery at West China Hospital (WCH). Disease stage was determined by the 8th edition of the American Joint Committee on Cancer (AJCC) TNM stage system (Goldstraw*et al*., 2016). This study was approved by the Institutional Review Board of West China Hospital of Sichuan University (Chengdu China; Project identification code: 2017.114 and 2018.270) and all patients signed informed written consent. Clinical characteristics including age, sex, smoking status, pathological subtype and stage were recorded at recruitment listed in Supplementary Table S1.

#### Quality check

Embedding samples with OCT: trim the sample to a suitable size, put it in an embedding box containing OCT embedding agent precooled on dry ice, quickly transfer the embedded samples to a refrigerator at −80°C on dry ice for quick freezing, and store them in the refrigerator at −80°C for a long time.

The OCT on the tissue surface was cut off, and the approximate tissue structure was seen. Trim the tissue surface in the cryomicrotome, extract 10 consecutive slices into the tube for RNA extraction, and post 3 consecutive slices on the slide for HE staining in the later stage

Stain processing.After the sample is activated, isopropanol acts for 1 minute to dry the sample until there is no water, hematoxylin stains for 7 minutes, anti-blue reagent for 2 minutes, eosin working solution for 1 minute, and then it is cleaned and dried and observed under the microscope

#### Tissue processing and Visium data generation

OCT (TissueTek Sakura) embedded samples cryosectioned at −10 °C (TermoCryostar). placeing the Sections on chilled Visium Tissue Optimization Slides (catalog no. 3000394, 10x Genomics) and Visium Spatial Gene Expression Slides (catalog no. 2000233, 10x Genomics), and warming the back of the slide to stuck firmly. Tissue sections were then fixed for 30mins in chilled methanol and then stained for next 30mins. For tissue optimization experiments, fluorescent images were taken with a a TRITC filter (Excitation 542/20,Emission 620/52) using a 10X objective and 300 ms exposure time, 18mins was the optimal time to permeabilize lung cancer tissue .For gene expression samples, which was selected 18mins time based on tissue optimization time course experiments. Taken brightfield histology images using a 10X objective (Plan APO) on a Olympus iX83 (2048 x2048 pixels for TO and GE) and the stitched raw images together using cellsens link (Olympus) and exported as .Tiff files with high resolution settings. According to the Visium Spatial Gene Expression User Guide Libraries were loaded at 300pM and sequenced on a NovaSeq 6000 System (Illumina) using a NovaSeq S1 Reagent Kit (200 cycles, 20028318, Illumina), at a sequencing depth of approximately 200 M read-pairs per sample. Sequencing was performed using the following theprotocolread 1, 51 cycles; i7 index read, 10 cycles; i5 index read, 10 cycles; read 2, 151 cycles.

#### Visium raw data processing

Raw FASTQ files and histology images were processed by sample with the Space Ranger software, which uses STAR v.2.5.1b (Dobin et al., 2013) for genome alignment, against the Cell Ranger hg38 reference genome “refdata-cellranger-GRCh38-3.0.0”, available at http://cf.10xgenomics.com/supp/cell-exp/refdata-cellranger-GRCh38-3.0.0.tar.gz.

#### Spatial Transcriptomics Data Processing

The gene-spot matrices generated after ST data processing from ST and Visium samples were analyzed with the Seurat package (versions 3.0.0/3.1.3) in R. In addition to custom scripts. For each patient data, spots were filtered for minimum detected gene count of 200 genes while genes with fewer than 10 read counts or expressed in fewer than 2 spots were removed, and Spots that account for more than 10% of mitochondrial genes are filtered. Normalization across spots was performed with the SCTransform function with regression of replicate and number of genes per spot. Dimensionality reduction and clustering was performed with Principal Components Analysis (PCA) at resolution 0.8 with the first 30 ICs. The spatial cluster genes differentially expressed (average logFC> 0.25 and adjusted p value < 0.05 by Wilcoxon Rank Sum test) across all ST clusters for each patient using FindAllMarkers function. Spatial feature expression plots were generated with the SpatialFeaturePlot function in Seurat (version 3.1.3)

#### Spatial Subclones Analysis

EMT gene set contain 13 mesenchymal marker genes (*VIM, CDH2, FOXC2, SNAI1, SNAI2, TWIST1, FN1, ITGB6, MMP2, MMP3, MMP9, SOX10, GCS*) and three epithelial marker genes (*CDH1, DSP, OCLN*), which reported important for tumor metastasis (Dev Dyn et al., 2018). Hallmark EMT signature scoring was performed using the AddModuleScore function in Seurat with default parameters. The immune score and tumor purity predict by ESTIMATE algorithm, which used gene expression data predicts the presence of stromal cells and immune cells in tumor tissues.pathway enrichment analysis were conducted with ClusterProfiler. Significance test uses Pearson Correlation Test for statistics (P<0.05).

#### Pattern of transient gene expression along spatial trajectory analysis

Pattern of transient gene expression along spatial trajectories was analyzed by SPATA, with (npcs = 30), FindNeighbors (dims = 1:20) for data preprocessing. By using assessTrajectoryTrends and filterTrends to model the spatial gene expression pattern, During the trajectory the expression was performed with the heatmap function in the ggplots package in R.

## Author Contributions

LZ conceived the project and designed the experiments., MLY, carried out experiments, SQM performed bioinformatic analysis. SQM, MLY, L.Z. drafted the manuscript. YY, NCC, TTS, YYN, YFY and ZQL contributed to the experiments and analyzed the data. All authors discussed the results and reviewed the manuscript.

## Supplementary Figure Legends

**Figure S1. A spatial visualization of the proportions of gene expression topics**. The Clinical Information in the survey. Interpolated tissue images how four gene expression topics with clear morphological patterns. Gene expression topic proportions are sample specific, meaning that the same proportion in another sample does not necessarily represent a similar region.

**Figure S2. The landscape of CNVs of subclones in the LUSC**. The landscapes of CNVs in LUSC, CNVs of subclones infer with inferCNV by using spatial transcriptome data. a. chosing the normal tissue with stromal cells (reference cells, top) as a control, calculated the copy number variation of tumor-related cluster 0,1,2,3 (observation cells), and the whole genome copy number results are presented through heatmap. a. The inferCNV result of subclones of LUSC_1, the CNVs score of four subclones had been shown in all of chromosomes. b. The inferCNV result of subclones of LUSC_1.

**Figure S3. The Intratumoral subclones heterogeneity and immune microenvironment heterogeneity**.

a, The specific and highly expressed genes of each of the 4 subclones, we selected the top two genes for display with Violin illustration. b. The Analysis of cluster 0 highly expressed gene GESA analysis shows that Hallmark Epithelial Mesenchymal Transition pathways are significantly enriched. c &d. Using ESTIMATE to calculation each subclones tumor microenvironmental components, this software can predict the purity of tumor. It is important to predict the purity of the tumor by predicting the immune score and stromal score, thereby predicting the content of stromal cells and immune cells. The LUSC_1 &LUSC_3 tumor purity and stromalscore, ImmuneScore had been shown and calculated the correlation between each score.

