## Supplementary material for "Spatial transcriptome sequencing revealed spatial trajectory in the Non-Small Cell Lung Carcinoma": Figure S1

a

| The Clinical Information | | | | | | | |
|---|---|---|---|---|---|---|---|
| ID | Gender | Age | Smoking | Differentiation | Pathological subtype | Clincial staging | TNM staging |
| LUAD_1 | F | 56 | Y | G2/G1 | LUAD | IVA | / |
| LUAD_2 | F | 68 | Y | G4 | LUAD | IB | / |
| LUAD_3 | M | 49 | Y | G1 | LUAD | IB | / |
| LUAD_4 | F | 51 | N | G4 | LUAD | IIIB | T3N2M0 |
| LUSC_1 | F | 66 | N | / | LUSC | IA | / |
| LUSC_2 | M | 69 | Y | G4 | LUSC | IIIA | pT3N1M0 |
| LUSC_3 | M | 51 | Y | / | LUSC | IIIB | T3N2M0 |
| LUSC_4 | M | 75 | Y | G4 | LUSC | IIA | T2bN0M0 |
| LUSC_5 | M | 53 | Y | G4 | LUSC | IIA | T2bN0M0 |
| LUSC_6 | M | 68 | Y | G4 | LUSC | IIB | pT3N0M0 |
| LUSC_7 | M | 39 | N | G3/G4 | LUSC | IIIA | T3N1M0 |
| LUSC_8 | M | 48 | N | / | LUSC | IIIB | T3N1M0 |

b

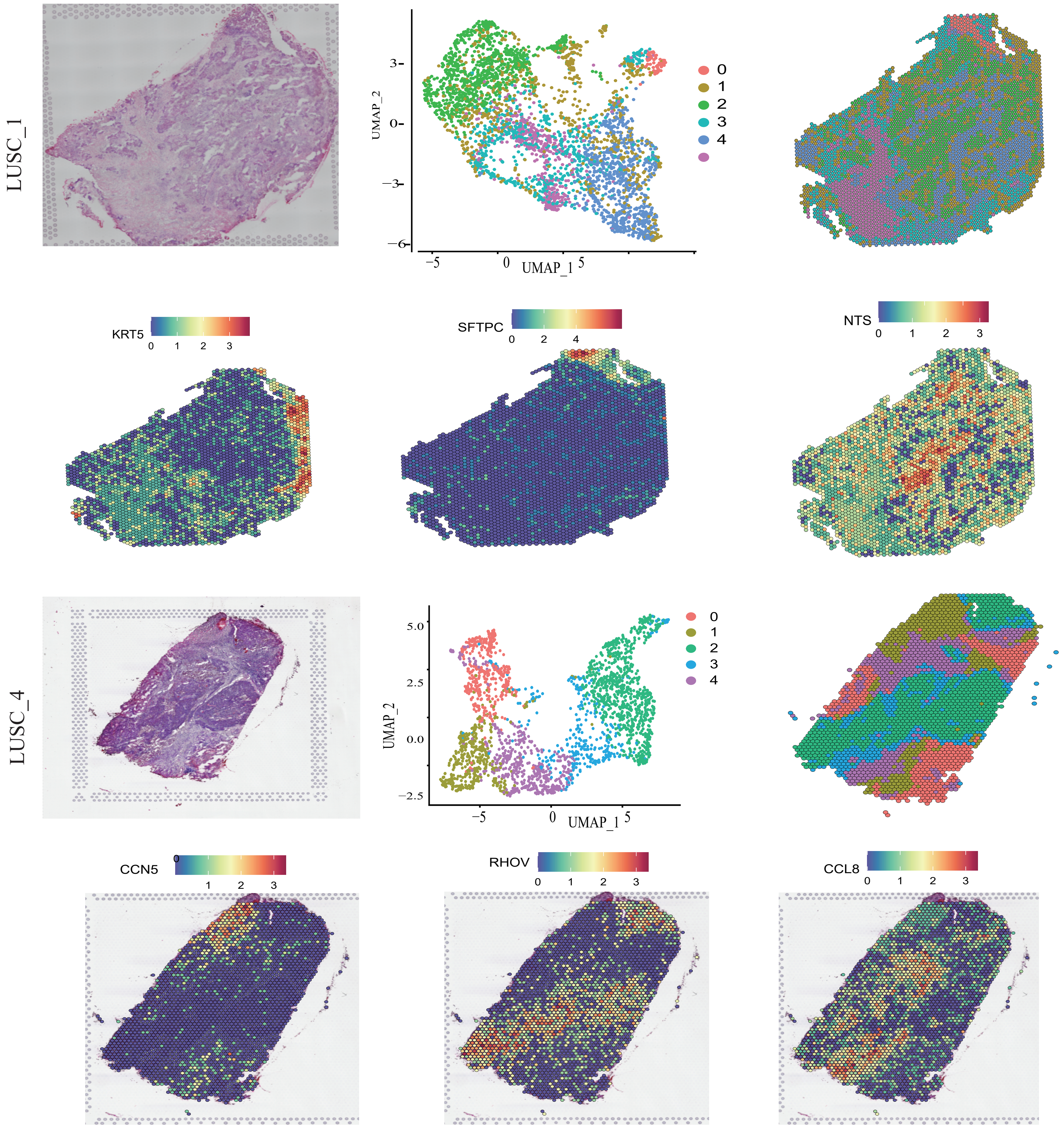
