## Supplementary figures and images for "Spatial transcriptome sequencing revealed spatial trajectory in the Non-Small Cell Lung Carcinoma"

### Figure S2

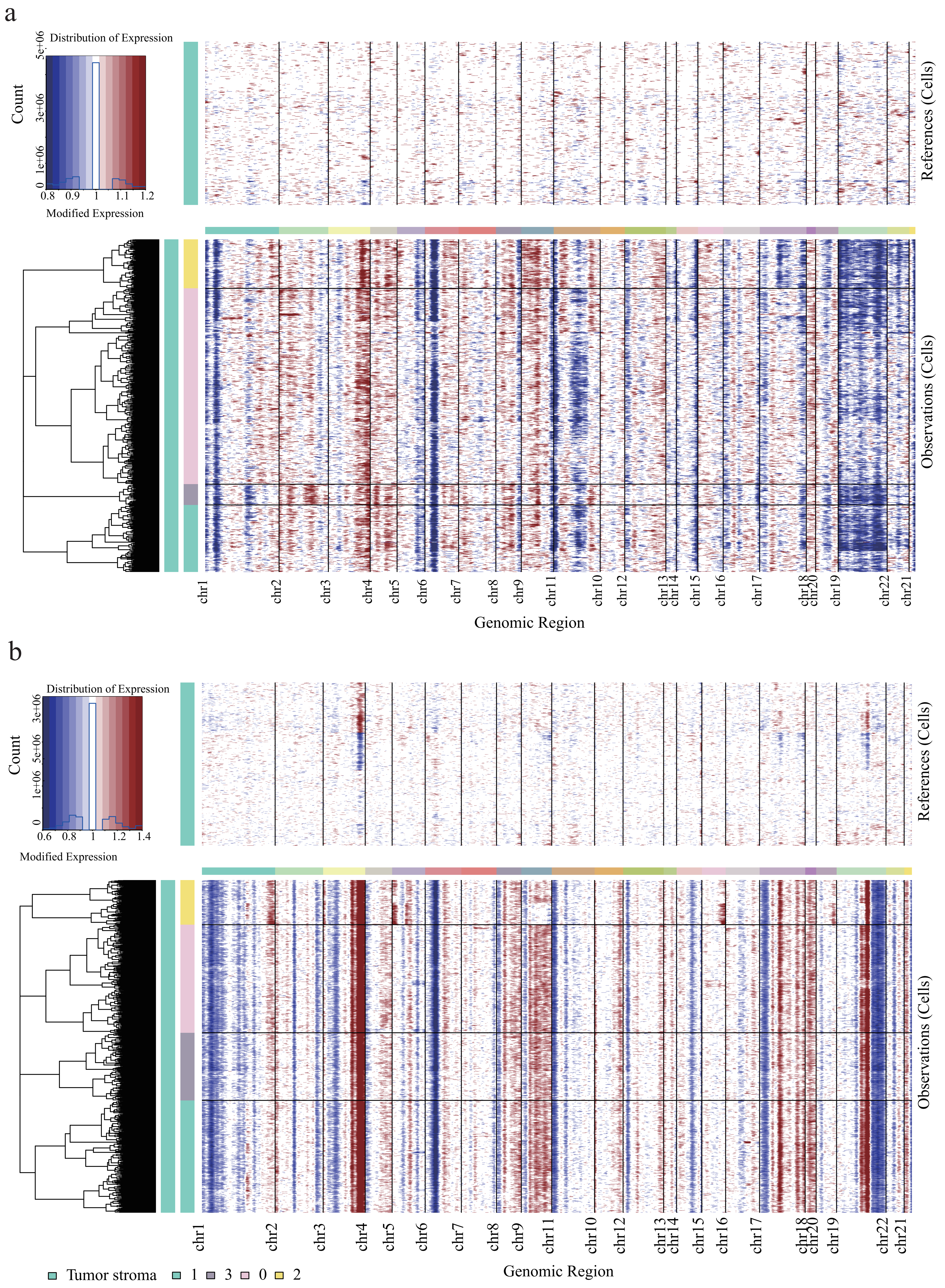

### Figure S3

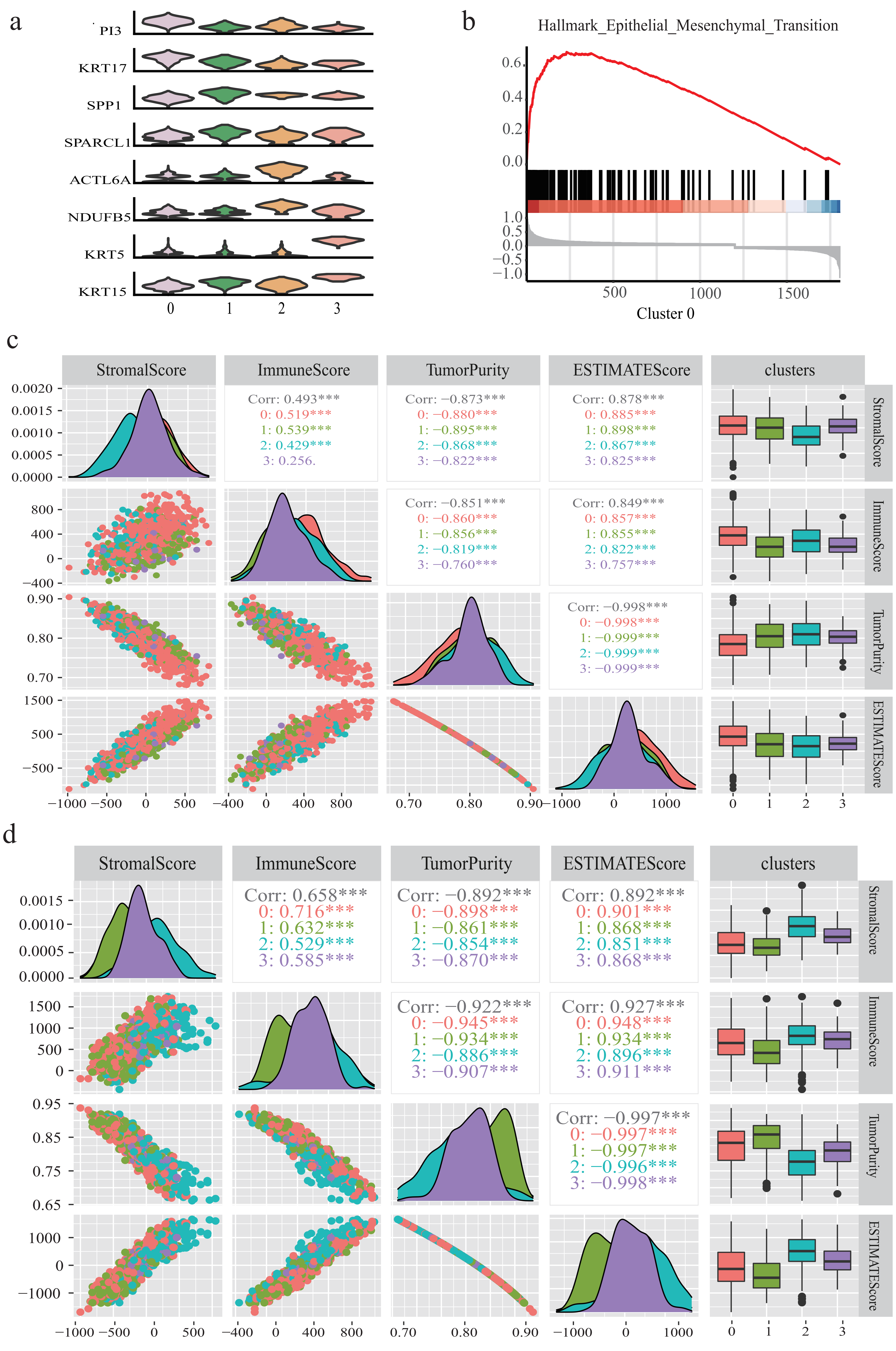
